# Pyocyanin produced by *Pseudomonas aeruginosa* Creates Legacy Effects That Boost Antibiotic Resistance Evolution in Enterococci

**DOI:** 10.64898/2025.12.04.692361

**Authors:** M.G.J. de Vos, V. Jansen, O. Bouhlali, A. Vlasblom, L.E. Zandbergen, I. van der Windt, J. Kool, R. Nijland, A. de Jong, O.P. Kuipers, S. Dunn, A. McNally, J.A.G.M. de Visser

## Abstract

Polymicrobial infections are small communities of multiple interacting bacterial species. Interactions among constituent species may modify the growth of community members in the presence of antibiotics, for example via degradation of the antibiotic or induction of specific resistance mechanisms. However, for most polymicrobial infections the nature of such interactions is opaque, while they may affect both treatment efficacy and the evolution of antibiotic resistance. Here, we describe that past interaction of enterococci with *Pseudomonas aeruginosa* creates legacy effects that substantially alter their antibiotic tolerance and resistance evolution. Specifically, we find that the temporary exposure to pyocyanin, a secondary metabolite produced by *P. aeruginosa*, increases the efflux in enterococci. These tolerance legacy effects promote the evolution of antibiotic resistance of enterococci. This work shows that transient interactions in polymicrobial communities can alter the evolutionary fate of community members.

## Introduction

Bacterial infections may consist of different species that together form a polymicrobial community, as is often the case in urinary tract infections (UTIs) ^1,2^. The response to antibiotic treatment of such polymicrobial infections may be affected by metabolic interactions among community members, that affect the survival and resistance evolution of specific members ^3–6^. For example, β-lactamase-producing strains may cross-protect community members against β-lactams ^7,8^ and antibiotic-resistant *Escherichia coli* strains producing the signal molecule indole may induce antibiotic tolerance via enhanced efflux in susceptible bacteria ^9^. Previous research of polymicrobial UTIs has shown that interactions between UTI members can alter the growth and antibiotic tolerance of other community members ^10,11^. Yet the nature of these interactions is often unknown.

The role of Gram-positive bacteria in infections has been debated, but evidence supporting the importance of these species in the polymicrobial UTI ecosystems is growing ^10,12–15^. Furthermore, the increased emergence and spread of antimicrobial resistance in these species ^16–18^ warrants investigation of their evolvability of antimicrobial resistance.

Here, we investigate the effect of interactions, with a panel of bacterial species that are frequently isolated together in persons diagnosed with polymicrobial UTIs ^1^, on the evolvability of antimicrobial resistance in enterococci.

## Results

### *P. aeruginosa* increases the production of antibiotic-resistant colonies in *E. faecium*

To test for cross-species interactions that would enhance antibiotic resistance in *Enterococcus faecium* – either through increased mutation rates, antibiotic tolerance or both, we investigated effects on the production of antibiotic resistant colonies in *E. faecium* from exposure to conditioned medium produced by *Escherichia coli, Klebsiella pneumoniae, Proteus mirabilis, Pseudomonas aeruginosa* and *Staphylococcus aureus*, species that often co-occur in polymicrobial UTIs. Details of the assay are described in the Material and Methods. Briefly, conditioned medium was generated by growing single isolates for 48 hours in liquid LB medium, recovering the supernatant, and replenishing it with nutrients. To screen for the differential evolution of resistance of *E. faecium* under the different conditioned media we adapted the P0-method (Luria Delbruck, 1943). We exposed 288-replicate 200 µL populations of *E. faecium* overnight to either conditioned medium of bacterial isolates or LB reference medium, and spotted 8-µL samples of each replicate overnight culture onto fresh LB agar plates (without conditioned medium) containing the antibiotic rifampicin (15 µg/mL, twice its minimal inhibitory concentration, MIC). Based on the number of replicate cultures that showed at least one antibiotic resistant colony, we calculated the evolvability (Materials and Methods). We define relative evolvability of antibiotic resistance in *E. faecium*, A, as the evolvability of antibiotic resistance in *E. faecium* exposed to conditioned versus unconditioned reference medium (LB): A = *a*_conditioned medium_*/a*_reference medium_. This method allows us to investigate the generation of resistant mutants in the different conditions, in the absence of selection for antibiotic resistance.

We find that the transient, overnight, exposure to most conditioned media (i.e., from *E. coli, K. pneumoniae, P. mirabilis*, and *S. aureus*) does not affect the evolvability of rifampicin resistance of *E. faecium*. However, there is one notable exception: the presence of the conditioned medium of *P. aeruginosa* increases evolvability ∼10 times compared to the populations that were not exposed to this conditioned medium (One-way ANOVA, *p* << 0.01, post-hoc Dunnett’s test *p* << 0.01, Figure 1A). Overnight regrowth of single colonies that appeared in the evolvability assay in LB (in the absence of antibiotics), followed by plating on 50 µg/mL rifampicin containing LB agar plates, showed that colonies were highly resistant (over 50 µg/mL rifampicin) and that resistance was heritable (Materials and Methods).

**Figure 1:**
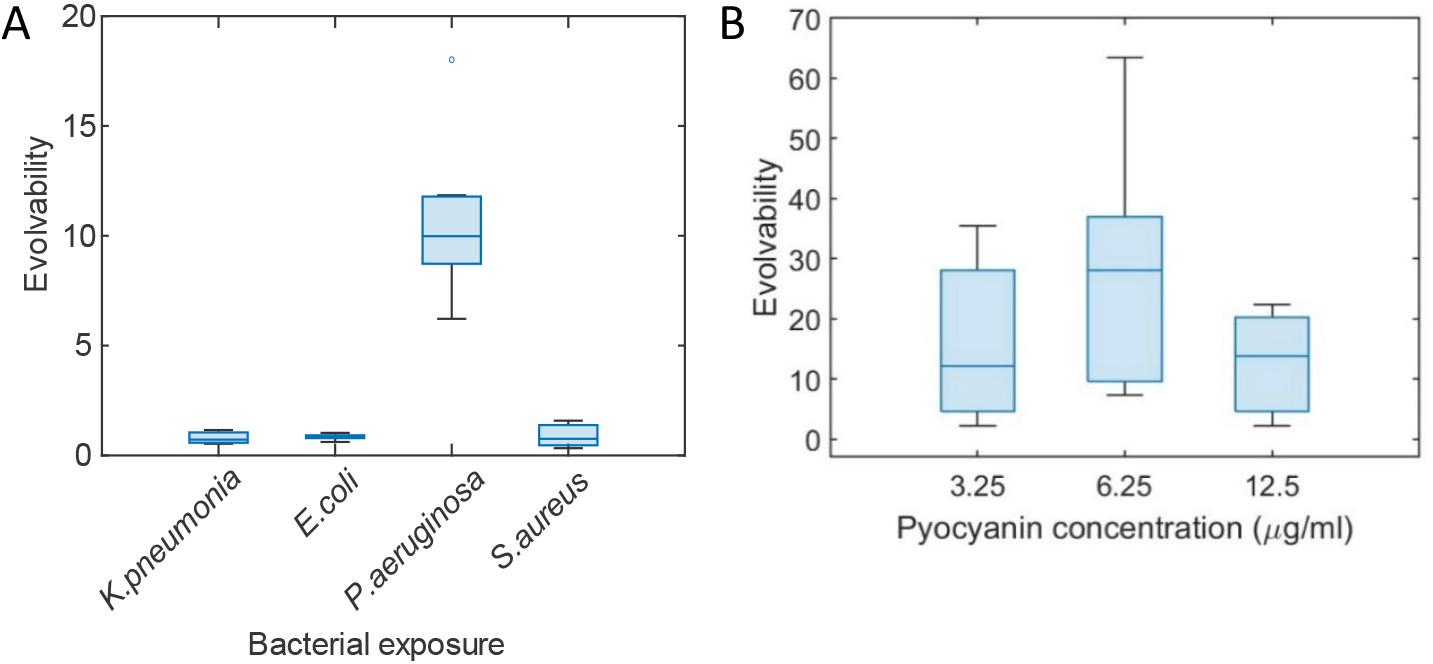
Evolvability of antibiotic resistance mediated by *P. aeruginosa* secondary metabolite. **A Evolvability rifampicin resistance of *E. faecium* under the influence of conditioned media derived from different bacterial species**. Evolvability of rifampicin resistance is measured as the rate of rifampicin resistance in conditioned medium normalized to the rate of rifampicin resistance in LB reference medium, as determined by the evolvability assay (Materials and Methods). The presence of the conditioned medium of *P. aeruginosa* increases the evolvability of rifampicin resistance of *E. faecium* (One-way ANOVA, *p* = 4.6e-22, post-hoc Dunnett’s test *P* = 1.2e-06 for *P. aeruginosa medium*). Other conditioned media of *K. pneumoniae, E. coli* and *S. aureus* do not alter the evolvability of antibiotic resistance. The lines in the box plots represent the median, bottom and top edges represent the lower and upper quartiles, whiskers the end points of the dataset, and small circle an outlier. **B Pyocyanin leads to increased evolvability of rifampicin resistance in *E. faecium***. Different physiologically relevant concentrations of pyocyanin (3.125, 6.25, 12.5 mg/mL) in LB lead to the increased evolvability of rifampicin resistance, as determined by the evolvability assay, normalized to the rate of rifampicin resistance in LB medium (One-way ANOVA, *p* =0.01, post-hoc Dunnet’s test, *p* =0.04 for 3.25 mg/mL, *p* =0.01 6.25 mg/mL and *p* = 0.4 12.5 mg/mL pyocyanin). The lines in the box plots represent the median, bottom and top edges represent the lower and upper quartiles, whiskers the end points of the dataset.

We know that conditioned media can alter the nutrient concentration and the constituents of the medium. We found no effect of the nutrient concentration on evolvability of resistance (Figure S1A). Testing a larger set of bacterial interactions via conditioned media (i.e. from two *E. coli* isolates, two *P. mirabilis* isolates, and single isolates of *S. haemolyticus, E. faecium* and *E. faecalis*), we find that the magnitude of the effect on evolvability of *E. faecium* is not related to population size as a consequence of the nutrient status of the conditioned medium, Figure S1B), in contrasts to findings of Kraṧovec et al, Krašovec et al., 2017). Conditioned medium can also lower the general efficacy of antibiotics, via other changes to the medium related to nutrient composition, for instance by increasing the pH ^10^. If the conditioned medium of *P. aeruginosa* had such an efficacy-altering effect on rifampicin, we would expect this to also affect other species than *E. faecium*. Therefore, to rule out that *P. aeruginosa*-conditioned medium altered the general efficacy of rifampicin, we performed the antibiotic resistance evolvability assay with two other focal species, *E. coli* and *K. pneumoniae. E. coli* and *K. pneumoniae* did not experience an increased phenotypic evolvability in the presence of *P. aeruginosa*-conditioned medium (one-way ANOVA, *p* = 0.25). We can, therefore, conclude that the increased evolvability of rifampicin resistance in *E. faecium* due to *P. aeruginosa*-conditioned medium is not an artifact caused by the decreased efficacy of rifampicin.

To test the generality of the increased antibiotic resistance evolvability of *E. faecium* after transient exposure to *P. aeruginosa*-conditioned medium, we performed a similar evolvability assay with nalidixic acid instead of rifampicin. Nalidixic acid has another mode of action than rifampicin, where rifampicin mainly inhibits RpoB, the RNA polymerase, nalidixic acid inhibits DNA gyrase activity ^20^. We find that the transient exposure to *P. aeruginosa*-conditioned medium also leads to the increased evolvability of nalidixic resistance of *E. faecium*, whereas the conditioned medium of *E. faecium* does not have this effect (one-way ANOVA, *p* = 0.0005, post-hoc Dunnett’s test, *p* = 0.0005 for *P. aeruginosa* conditioned medium, *p* = 0.96 for *E. faecium* conditioned medium, Figure S2). These combined results suggest a particular interaction of *P. aeruginosa* and *E. faecium* on the evolvability of antibiotic resistance.

### Genetic basis of resistance

To further test whether the increased evolvability of *E. faecium* due to the transient exposure to *P. aeruginosa*-conditioned medium corresponded to an increased genomic mutation rate, we sequenced the genomes of evolved isolates derived from the phenotypic evolvability assay. By means of whole genome sequencing of highly resistant (> 50 µg/ml rifampicin) mutants that appeared on the rifampicin (15 µg/ml) containing agar plates, we found that mutants derived after the transient exposure to *P. aeruginosa* conditioned medium had on average 50% more, as well as different, mutations compared to those exposed to LB (Table S1 and S2, Figure S3). Yet the difference in the distribution of mutant numbers was not statistically significant (Student’s t-test, *p* = 0.09). Interestingly, clones exposed to LB medium contained canonical rifampicin resistance *rpoB* mutations ^21^, whereas the clones exposed to *P. aeruginosa* conditioned medium mostly contained different *rpoB* mutations (Table S1). Moreover, clones derived from the conditioned medium of *P. aeruginosa* contained other additional genomic mutations, outside of *rpoB*, such as multidrug resistance protein *stp* ^22^, *pyrF* ^23^ *nrnA* ^24^, which are involved in the production and degradation of nucleotides ^25^ (Table S2).

The slight increase in the number of mutations in the genomes of resistant mutants after the transient exposure to *P. aeruginosa* conditioned medium suggests that DNA protection and repair proteins of stress response pathways may be involved. Indeed, the transcriptomic analyses reveal a down-regulation in *P. aeruginosa* relative to *E. faecium*-conditioned medium of *dps* (DNA protection during starvation protein, Log2-fold change −2.73, FDR 8.78e-3, p-value 5.73e-5), which is involved in oxidative stress protection and has ferroxidase activity, and *usp* ^*26*^ (universal stress protein, Log2-fold change −2.60, FDR 9.47e-4, p-value 2.20e-5), involved in general stress protection. We did not find evidence for the involvement of other known stress response genes involved in the SOS or stringent response. While the downregulation of protection and repair genes may point to the mechanism behind the 50% increase in number of mutations per clone, this clearly cannot account for the >10-fold upregulation of evolvability. This suggests that another factor must play a role in the increased phenotypic evolvability of antibiotic resistance.

### *P. aeruginosa* induces lasting antibiotic tolerance

To investigate whether the observed increased evolvability is (partly) due to the physiological induction of certain resistance mechanisms, we performed an antibiotic tolerance assay. We grew *E. faecium* in a two-fold dilution gradient of rifampicin and nalidixic acid, in the presence and absence of *P. aeruginosa*-conditioned medium. We find that *P. aeruginosa*-conditioned medium indeed increases the tolerance to these antibiotics (Figure S4), by allowing *E. faecium* to grow at higher concentrations of antibiotic compared to in LB reference medium. This antibiotic tolerance effect, the decreased sensitivity to antibiotics, is in line with the previously observed induced tolerance of nitrofurantoin and sulfamethoxazole-trimethoprim combination ^10^, and vancomycin ^6^ of enterococci in the presence of *P. aeruginosa*.

To investigate whether the effect of the transient, overnight, exposure to *P. aeruginosa*-conditioned medium that we describe above is short-lived, we repeated the evolvability assay with two different additional washing steps, before spotting the cultures on the rifampicin-containing LB agar plates. Specifically, we washed a subset of the overnight cultures exposed to *P. aeruginosa*-conditioned medium in PBS before spotting the replicate cultures on the rifampicin-containing LB agar plate. Additionally, we washed a subset of overnight-exposed cultures in LB followed by a three-hour regrowth in LB (in the absence of conditioned medium), before spotting samples on the rifampicin-containing agar plates. Neither washing with PBS, nor a three-hour regrowth in fresh LB, did ameliorate the increased phenotypic evolvability due to the exposure to *P. aeruginosa* conditioned medium (Figure S5A). This shows that the increased evolvability induced by *P. aeruginosa* conditioned medium is rather long lasting.

### *P. aeruginosa* increases antibiotic resistance in enterococci

To investigate whether other isolates of *P. aeruginosa* and other enterococci also experienced increased phenotypic antibiotic resistance, we tested the effect of canonical *P. aeruginosa* isolate PAO1, and the uropathogenic *P. aeruginosa* isolate used above, on three uropathogenic *E. faecalis* strains from our collection ^1^. These followed a similar pattern, with for some combinations an even higher elevated phenotypic evolvability of resistance, induced by the transient overnight exposure to *P. aeruginosa*-conditioned medium (Figure S5B). This suggests that the increased evolvability is not limited to these initially tested isolates, but is rather due to a general *P. aeruginosa* – enterococci interaction.

### Pyocyanin produced by *P. aeruginosa* induces increased efflux in enterococci

We noted that particularly the PAO1-conditioned medium with a green tint, but not the un-tinted PAO1-conditioned medium or un-tinted PAO1 quorum-sensing-mutant conditioned medium generated under lower oxygen conditions, led to a higher number of resistant colonies, and therefore a strongly increased the antibiotic resistance evolvability of *E. faecium* (Figure S6A). This led us to hypothesize that pyocyanin, which is responsible for the colored hue in the conditioned medium, is produced under aerobic conditions and plays an important role in the virulence of *P. aeruginosa* ^27,28^, causes the increased numbers of rifampicin-resistant colonies in the evolvability assay. Testing the effect of physiologically relevant concentrations of pyocyanin ^29^on the evolvability of rifampicin resistance indeed showed an increase of rifampicin-resistant colonies (Figure 1B). Phenazine, a structurally similar compound and precursor of pyocyanin, did not lead to an increase in rifampicin resistance (Figure S6B).

To investigate the mechanistic basis underlying this lasting, legacy effect on the evolvability of antibiotic resistance, we compared the gene expression levels of *E. faecium* exposed to *E. faecium* versus *P. aeruginosa*-conditioned medium. The results showed that under the influence of *P. aeruginosa* several efflux pumps and transport systems displayed increased expression (Figure 2A, Figure S7, Table S3). Particularly, ABC transporter MacB Fts-X-like permease is upregulated. Genes in this complex have previously been implicated to be involved in drug tolerance via the efflux of antibiotics, particularly macrolides and cephalosporins, but also colistin and bacitracin ^30,31^.

**Figure 2:**
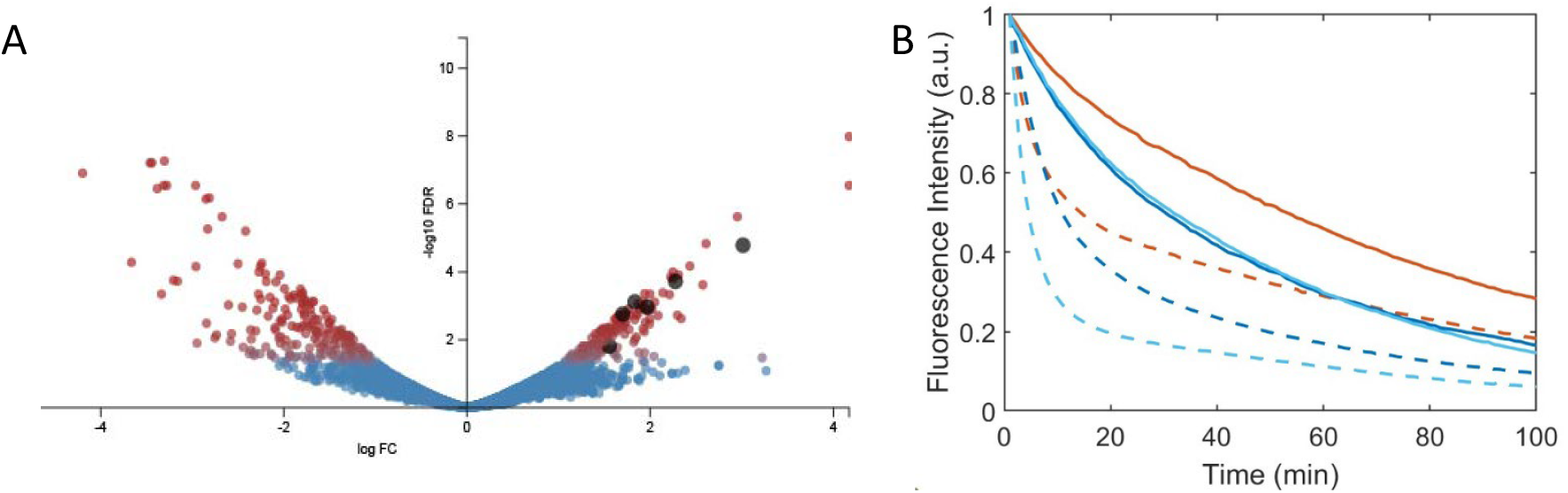
Efflux upregulated in *Enterococcus faecium* exposed to *P. aeruginosa* conditioned medium. **A Differential gene expression analysis by means of transcriptomics of *E. faecium* cultured under *P. aeruginosa* versus *E. faecium* conditioned medium shows the upregulation of several transporters**. Red dots represent significantly differently expressed genes, defined as ≥1.5 log_2_ fold change (FC), and <0.05 false discovery rate (FDR). Several transporters (black dots) are upregulated in the presence of conditioned medium of *P. aeruginosa* compared to the presence of *E. faecium* (control) conditioned medium. The most highly upregulated transporter is denoted as ABC transporter MacB Fts-X like permease. **B Ethidium bromide efflux assay of *E. faecium* in the presence of *P. aeruginosa* conditioned medium, pyocyanin and LB control medium**. Overnight exposure of *E. faecium* to *P. aeruginosa* conditioned medium (dark blue solid line), and pyocyanin (5 ug/ml, light blue solid line) increase efflux in *E. faecium* compared to the LB reference medium (red solid line). In the presence of glucose (dashed lines) efflux is further enhanced, because efflux is an energy-requiring process.

To investigate whether the increased gene expression of efflux-related transporters translated in an upregulated efflux phenotype, we performed an efflux assay in which we exposed *E. faecium* cells to either *P. aeruginosa* conditioned medium, pyocyanin or reference LB medium. This showed that overnight exposure to the conditioned media of *P. aeruginosa*, as well as to a physiological relevant concentration of pyocyanin in LB, leads to increased efflux relative to reference LB medium (Figure 2B). Therefore, we conclude that pyocyanin produced by *P. aeruginosa* leads to increased efflux and increases tolerance to antibiotics.

The increased antibiotic tolerance and survival creates potential for increased evolvability via the selection of mutations causing antibiotic resistance ^32,33^. The upregulation of efflux-pumps and concomitant down-regulation of DNA repair and protection mechanisms, has previously been implicated to lead to increased evolution of antibiotic resistance evolution ^34^, which explains the higher numbers of rifampicin-tolerant *E. faecium* colonies in the evolvability assay after the transient exposure to *P. aeruginosa*-condition medium (Fig. 1). Together, this indicates that the transient exposure of enterococci to conditioned medium of *P. aeruginosa* containing pyocyanin leads to the increased tolerance due to increased efflux of antibiotics, which in turn leads to the increased evolvability of antimicrobial resistance.

## Discussion

Here, we show that the transient exposure of the Gram-positive enterococci to Gram-negative *P. aeruginosa*-conditioned medium leads to the increased evolvability of antimicrobial resistance, specifically rifampicin and nalidixic acid, which is correlated with the down-regulation of DNA repair enzymes and the increased efflux induced by pyocyanin, a secondary metabolite produced by *P. aeruginosa*. Overexpression of efflux pump genes has been associated with antibiotic resistance in various species ^35–37^. Additionally, the increased expression of efflux has been associated with the relatively fast evolution of antibiotic resistance^38^ ABC transporters such as MacB-FtSX-like permease have previously been implicated in increased resistance towards antibiotics, such as vancomycin ^39^. We show that efflux can be upregulated due to the presence of interspecies signals, such as pyocyanin.

Pyocyanin has previously been implicated in the induced tolerance to several antibiotics in *P. aeruginosa* ^40^, due to the internal intracellular accumulation of pyocyanin. Additionally, pyocyanin has been shown to increase SoxR mediated efflux in *P. aeruginosa* ^*41*^. Here, we report the interspecies signaling effect of pyocyanin, leading to increased efflux activity, and concomitant reduced sensitivity to antibiotics, such as structurally dissimilar rifampicin and nalidixic acid, in Gram-positive enterococci. This interspecies signaling appears to be rather specific, because the evolvability of Gram-negative *E. coli* and *K. pneumonia* remained unaffected by the conditioned medium of *P. aeruginosa*. Previous studies also showed that pyocyanin affected the tolerance and evolution of resistance to structurally similar synthetic antibiotics in Gram-negative *Stenotrophomonas maltophilia* and *Burkholderia cepacian* complex ^42^. Additionally, the efflux-inducing effect of indole signaling among antibiotic-resistant and susceptible bacteria within *E. coli* populations was shown to affect tolerance to norfloxacin ^9^. We therefore anticipate that similar interactions, mediated by secondary metabolites, within and across bacterial species could be more widespread and may explain the mismatch between observed clinical antibiotic levels and *in vivo* treatment failure ^43,44^.

The increased evolvability of antibiotic resistance of enterococci due to the transient exposure to *P. aeruginosa*-conditioned medium was further shown to last after removal of the conditioned medium, indicating substantial bacterial memory ^45^. We hypothesize that the upregulation of efflux pumps leads to the production and accumulation of efflux proteins that is sufficiently high to have a lasting effect on antibiotic efflux, even after several cell divisions on agar in the absence of the efflux-inducing presence of pyocyanin. Under such circumstances, cells experience longer survival and larger population sizes in the presence of antibiotics, potentiating the production of antibiotic resistant mutants and high resistance levels.

Due to the general effect of conditioned medium of *P. aeruginosa* on enterococci, we propose that future work should investigate whether other secondary metabolites, similar to pyocyanin, have similar inter-species efflux-inducing properties promoting antimicrobial tolerance, also in other Gram-positive species. Additionally, understanding the molecular mechanism of induction of efflux pumps, and the downregulation of DNA repair mechanisms, is mandatory to understand the scope of this cross-species effect on antibiotic tolerance.

Ours study shows, that not only current interactions, but also past metabolic interactions with other bacterial species can have a lasting effect on the growth of enterococci in environments containing antibiotics. Such lasting, legacy interactions have the potential to change the eco-evolutionary interactions in polymicrobial infection ecosystem formation related to antimicrobial resistance.

## Materials and Methods

### Isolates

We used isolates derived from polymicrobial urinary tract infections obtained for previous studies (Croxall et al., 2011), except for *P. aeruginosa* PAO1 and its derivative quorum-sensing mutants (LasR, LasI) which were a kind gift from Ashleigh Griffin (University of Oxford).

### Media

Liquid Luria Broth (LB) was created as follows: 10 g/L bacto tryptone, 5 g/L yeast extract; 10 g/L NaCl. 5x LB: 50 g/L bacto tryptone; 25 g/L yeast extract, no NaCl. Agar plates were made using LB with 15 g/L Technical agar #3 (Sigma), and different concentrations of antibiotics. Antibiotic testing and colony formation in evolvability assays were performed on 1xLB agar plates, containing rifampicin (15 μg/mL, 50 μg/mL, 100 μg/mL), or nalidixic acid (150 μg/mL). Refence medium to compare to conditioned medium in evolvability assays consisted of 1.5x LB including 0.01 M PIPES buffer pH 6.5.

### Conditioned medium

Donor strains were grown in 500 mL Erlenmeyers in 200 mL LB under aerobic conditions for 48 hours at 37 °C at 200 rpm. All cultures were divided over 50 mL tubes and centrifuged for 15 min at 4700 rpm at room temperature. Supernatant was collected, leaving the pellet as undisturbed as possible. The cultures were filtered using a 0.45 μm mesh filter attached to a vacuum pump set at 0.1-0.5 Bar. A 0.2 μm mesh filter was used to filter the cell-free supernatant for a second time. Conditioned medium was generated by mixing 0.5 fraction spent medium (cell-free supernatant), 0.1 fraction filter sterilized 0.01 M PIPES buffer, pH 6.5, 0.1 fraction 5x LB, 0.3 fraction 1xLB, such that in the hypothetical case that no nutrients would have been consumed in the conditioned medium the concentration of LB would be similar to the reference medium.

### Antibiotic resistance

The level of antibiotic resistance before performing the evolvability assays was assessed by spotting 10 μl of overnight culture on LB agar plates containing the appropriate antibiotics. The level of resistance was determined by the highest concentration at which growth was observed on the LB agar plate containing a certain concentration of antibiotic.

### Evolvability assay

The high-throughput evolvability assay is based on the fluctuation assay developed by Luria and Delbrück to estimate mutation rates of bacterial strains ^46^. One pre-culture was made in LB and diluted in pre-warmed LB to a density of 10^-3^. 23 µL of this diluted culture was added to 23 mL of LB, conditioned medium (inoculated density OD600 10^-6^), LB plus pyocyanin or LB plus phenazine. Per condition at least two 96-well plates were filled with 200 µL of bacteria medium mixture. These 96-well plates were incubated overnight, under non-shaking conditions at 37 °C. After 16 hours, the plates were placed in a BMG Clariostar plate reader, and mixed for 240 seconds at 600 rpm, double orbital, and the OD600 was recorded. Following mixing and OD600 measurement, 8 µL drops from each well were spotted on pre-warmed LB agar plates containing rifampicin or nalidixic acid (resp. 15 µg/ml and 150 µg/ml, i.e. twice the MIC concentration). For Figure 1, 288 replicate cultures, and for Figure S1, at least 192 replicate cultures in each condition were used to calculate the rate of resistance production. After two days of incubation at 37 °C, the number of spots that had at least one antibiotic-resistant colony, were counted. The evolvability measure was calculated using the p-0 method as:

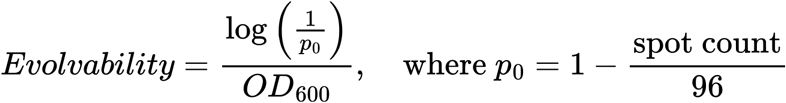

This calculation allows for normalization of the samples according to their population size after overnight growth estimated by the optical density at 600 nm (OD600). The evolvability of isolates exposed to conditioned medium was normalized to the evolvability of isolates exposed to LB.

### Assessment of resistance level of colonies derived from evolvability assay

The resistance level of the clones that appeared in the evolvability assay was assessed by regrowing clones overnight in LB and spotting ∼2 µL of this overnight culture on fresh agar plates containing rifampicin (50 µg/mL, 100 µg/mL). Isolates showing growth at the respective agar plates containing antibiotics were scored as resistant.

### Antibiotic tolerance assay

Two µL of *E. faecium* overnight cultures was reinoculated in 96-well plates in liquid cultures for 24 h. The effect of *P. aeruginosa* conditioned medium on the action of antibiotics was assessed by scoring the yield in a concentration gradient of rifampicin and nalidixic acid in the presence and absence of *P. aeruginosa*-conditioned medium, using LB as reference medium. Two-fold decrements and an additional no-drug control were used to determine the maximum antibiotic concentration that could sustain growth. Antibiotic tolerance interactions were scored as the difference in the number of antibiotic concentrations in which growth was observed in conditioned medium versus the LB reference. Growth of *E. faecium* was scored positive if OD600 exceeded 0.01 within 24 h. Antibiotics were added in the conditioned medium just before the start of the tolerance assay, i.e., they were absent during the growth of the donor from which the conditioned medium was produced.

### Ethidium bromide efflux assay

Five mL *E. faecium* cultures were grown for 20 hours at 37 °C in three conditions: in LB, conditioned medium of *P. aeruginosa*, or LB with 5 µg/mL pyocyanin (resolved in DMSO), with shaking at 200 rpm. After determination of the cell density (OD600), the cultures were spun down for 10 minutes at 4,500 rpm. Cultures were resolved in pre-warmed PBS at a density of 0.1 OD600. These samples were incubated in PBS in 15 mL tubes for 1 hour at 37 °C, in the presence of 5 µg/mL ethidium bromide. After incubation, cultures were spun down for 10 minutes at 4,500 rpm. The supernatant was discarded and 1 mL fresh PBS was added. The OD600 was adjusted to 0.1 and 200 µL of these suspensions (of *E. faecium* cultured under these three conditions) with and without 0.4% glucose (efflux is an active process requiring energy, energy is provided in the form of glucose) were added to a 96-well plate, with four replicates for each condition. Fluorescence development was monitored in a BMG Clariostar plate reader (excitation 520 nm, emission 590 nm).

### DNA extraction

Genomic DNA was isolated using a phenol/chloroform extraction based on the method of Sambrook ^47^ with some modifications as described. Bacterial cells were spun down and cell pellets were washed in 250 µL PBS, and resuspended in 250 µL PBS. Ten µL of 20 mg/mL lysozyme was added and incubated for 30 minutes at 37 °C. Fifteen µL of 10 % SDS and 5 µL of 20 mg/mL RNAseA were added, and incubation was continued at 37 °C for 30 minutes. Fifteen µL of ProtK (20 mg/mL) was added, and the sample was further incubated at 56 °C for 30 minutes. 300µL of Phenol:Chloroform:Isoamyl alcohol (Sigma-Aldrich) was added, the sample was mixed on a rotator at 25 rpm for 15 minutes and centrifuged for 10 minutes at 14,000 rpm. Supernatant was collected, and DNA was precipitated by the addition of 10% 3M NaAcetate and 1 mL 100% ice-cold ethanol. DNA was pelleted by centrifugation at 10,000 rpm for 5 minutes, and the pellet was washed twice with 300 µL 70% ethanol. The pellet was shortly dried and dissolved in 50 µL nuclease free water. To remove residual RNA and further purify the DNA, a second RNAse treatment and double P:C:IAA clean-up was performed as follows. 30 µL of DNA solution was diluted by addition of 150 µL of Tris-EDTA buffer and 2 µL if 20 mg/mL RNaseA. Samples were incubated at 37 °C for 1 hour. 250 µLof Phenol:Chloroform:Isoamyl alcohol was added, mixed and centrifuged as describe above. 200 µL of the supernatant was transferred to a 2 mL phase-lock tube, 200 µL of Phenol:Chloroform:Isoamyl alcohol was added, mixed and centrifuged again as described above. 180 µL supernatant was transferred to a 1.5 mL DNA lowbind tube (Eppendorf) and 10% 3 M NaAcetate (pH 5.5) and 800 µL icecold 80% ethanol was added, and the tube was incubated on ice for 1 h to precipitate the DNA. DNA was collected by pelleting as described above, washed twice in 300 µL 70% ethanol, airdried at room temperature for 2 minutes and dissolved in 30 µL nuclease free water.

### Whole genome sequence analysis

The parental *E. faecium* strain was sequenced on the Illumina Novaseq and ONT Minion platform by MicrobesNG (https://microbesng.com) to create a closed reference genome sequence, assembled using Unicycler and annotated using Prokka. Evolved strains were also sequenced on the Illumina Novaseq platform by MicrobesNG and reads filtered using TrimAdapt. The resulting FastQ files were then mapped against the closed parental strain genome using BreSeq.

### RNA extraction, library prep and transcriptomic analysis

*E. faecium* cultures for RNA extraction were prepared by creating an overnight culture in LB broth with growth at 37 °C, shaken at 200 rpm. This overnight culture was used to inoculate the different conditioned media and *E. faecium* was allowed to grow halfway into its exponential phase (OD ≈ 0.15) at 37 °C, 200 rpm. Cultures were pelleted at 4,700 rpm for 15 minutes and supernatants were discarded. Pellets were suspended in 0.5 mL medium and transferred to Eppendorf tubes and centrifuged again at 18,000 rpm for 5 minutes. After this, the supernatants were discarded and pellets were flash frozen in liquid nitrogen for RNA extraction.

RNA was isolated using High Pure RNA Isolation Kit (Roche), with an adapted protocol for improved RNA microarray analysis. Pellets were thawed on ice and resuspended in 400 μL DEPC-treated MQ (100 µL DEPC per 100 mL solution). This solution was added to screw-cap tubes containing 0.5 g glass beads, 50 µL 10% sodium dodecyl sulfate (SDS) and 500 µL acid-phenol:chloroform, pH 4.5 (with Indole-3-acetic acid (IAA), 125:24:1) (ThermoFisher AM9720). Tubes were placed in a bead beater (Mini-BeadBeater; 607, BioSpec, USA) and a 6 × 60 s pulse homogenization with 1 min interval on ice was performed. Tubes were subsequently centrifuged at 10,000 rpm for 10 minutes at 4 °C. The resulting upper phase was transferred to a fresh tube with 500 µL chloroform:IAA (24:1) and centrifuged again for 5 minutes at 10,000 rpm at 4°C. 500 µL of the upper phase was again transferred to a new tube with 1 mL of the Roche lysis/binding buffer (4.5 M guanidine HCl, 30% Trition-X-100, 50 mM Tris, pH 6.6) and mixed well. 800 µL was transferred to the upper reservoir of the provided filter and collection tube. Tubes were centrifuged for 15 seconds at 8,600 rpm. Flow through was discarded and the remainder was transferred back to the upper reservoir of the filter. The tube was centrifuged again for 15 seconds at 8,600 rpm. Flow through was discarded and 100 mL of DNaseI mix was added, containing 90 µL DNase buffer and 10 µL DNaseI. Tubes were incubated for 30 minutes at 25 °C. 500 µL wash buffer I was added and the tubes were centrifuged at 8,600 rpm for 15 seconds. This was repeated with wash buffer II, and again with 200 µL wash buffer II and centrifuged at max speed. The filter was then transferred to a sterile 1.5 mL Eppendorf tube. Fifty µL elution buffer was added directly on the filter in the upper reservoir and incubated for 10 minutes at room temperature. Tubes were centrifuged for 1 minute at 8,600 rpm and aliquots were stored at −80 °C until the library prep.

RNA concentrations were measured using a Nanodrop 2000c Spectrophotometer (Thermo Fisher Scientific, USA) and the quality of RNA was analyzed using gel electrophoresis. A 50 mL gel was created of 1% w/v agarose in TAE buffer with 500 µL Bleac (Bleac is created by dissolving 1 tablet of a RECA bleach tablet in 100 mL MilliQ water). The mixture was incubated at room temperature for 5 minutes before heating the agarose to boiling temperature. Agarose was allowed to cool to 65 °C before adding 2 mL ethidium bromide. After pouring the gel and letting it set, RNA samples were mixed with 2x RNA loading buffer. The 2x RNA loading buffer consisted of 9.5 mL formamide, 25 µL of 10% SDS, 2.5 mg bromophenol blue, 2.5 mg ethidium bromide, and 10 µL of 500 mM EDTA. The gel was run in TAE buffer at 100 V for 25 minutes.

RNA was reverse-transcribed into cDNA using the reagents from the Zymo-Seq Ribofree cDNA kit (Zymo Research). DNA was resuspended in 10 µL DNA elution buffer and the mixture was transferred to a new PCR tube to be stored at −20 °C before sending it for sequencing. Samples were sequenced on the Illumina NexSeq 500 to generate 75 bases single end reads (75SE) with an average read depth of 12 million reads per sample.

Differential gene expression was quantified using Kallisto (V 0.46.0) with read length set to 75 bp and standard deviation set to 5. The resulting data was analyzed using Voom/Limma in Degust (V 3.20) to identify significantly differentially transcribed genes (average Log fold change of ≥1.5, and a FDR significance value of <0.05). Functional categories (COG, GO-terms) were assigned to genes using eggnog-mapper (V 2).

## Acknowledgements

We kindly thank Ashleigh Griffin (University of Oxford) for sharing *P. aeruginosa* PAO1 and quorum sensing mutants. MGJDV was supported by an NWO VENI Fellowship (863.14.015) and L’Oreal Unesco LNVH NIAS-KNAW For Women in Science Fellowship, part of this work was performed at NIAS-KNAW. LEZ was supported by the FSE Adaptive Life Program.

## References

1. Croxall, G. et al. Increased human pathogenic potential of Escherichia coli from polymicrobial urinary tract infections in comparison to isolates from monomicrobial culture samples. J Med Microbiol 60, 102–109 (2011).

2. Neugent, M. L. et al. Recurrent urinary tract infection and estrogen shape the taxonomic ecology and function of the postmenopausal urogenital microbiome. Cell Rep Med 3, (2022).

3. Zampieri, M. et al. Metabolic constraints on the evolution of antibiotic resistance. Mol Syst Biol 13, 917 (2017).

4. Windels, E. M. et al. Antibiotic dose and nutrient availability differentially drive the evolution of antibiotic resistance and persistence. ISME J 18, (2024).

5. Laborda, P., Martínez, J. L. & Hernando-Amado, S. Evolution of Habitat-Dependent Antibiotic Resistance in Pseudomonas aeruginosa. Microbiol Spectr 10, (2022).

6. Zandbergen, L. E. et al. Microbial interactions affect the tempo and mode of antibiotic resistance evolution. bioRxiv 2024.06.06.597700 (2024) doi:10.1101/2024.06.06.597700.

7. Yurtsev, E. A., Chao, H. X., Datta, M. S., Artemova, T. & Gore, J. Bacterial cheating drives the population dynamics of cooperative antibiotic resistance plasmids. Mol Syst Biol 9, 683 (2013).

8. Zhao, X., Ruelens, P., Farr, A. D., de Visser, J.A.G.M. & Baraban, L. Population dynamics of cross-protection against β-lactam antibiotics in droplet microreactors. Front Microbiol 14, 1294790 (2023).

9. Lee, H. H., Molla, M. N., Cantor, C. R. & Collins, J. J. Bacterial charity work leads to population-wide resistance. Nature 467, 82–85 (2010).

10. de Vos, M. G. J., Zagorski, M., McNally, A. & Bollenbach, T. Interaction networks, ecological stability, and collective antibiotic tolerance in polymicrobial infections. Proceedings of the National Academy of Sciences 114, 10666–10671 (2017).

11. Lara, E. G., van der Windt, I., Molenaar, D., de Vos, M. G. J. & Melkonian, C. Using functional annotations to study pairwise interactions in urinary tract infection communities. Genes (Basel) 12, 1221 (2021).

12. Cruz, M. R., Graham, C. E., Gagliano, B. C., Lorenz, M. C. & Garsin, D. A. Enterococcus faecalis inhibits hyphal morphogenesis and virulence of Candida albicans. Infect Immun 81, 189–200 (2013).

13. Xu, W., Fang, Y. & Zhu, K. Enterococci facilitate polymicrobial infections. Trends Microbiol 0, (2023).

14. Kline, K. A. & Lewis, A. L. Gram-Positive uropathogens, polymicrobial urinary tract infection, and the emerging microbiota of the urinary tract. Microbiology Spectrum 4 (2016)

15. Keogh, D. et al. Enterococcal metabolite cues facilitate interspecies niche modulation and polymicrobial infection. Cell Host Microbe 20, 493–503 (2016).

16. Zaheer, R. et al. Surveillance of Enterococcus spp. reveals distinct species and antimicrobial resistance diversity across a One-Health continuum. Scientific Reports 2020 10:1 10, 1–16 (2020).

17. Markwart, R. et al. The rise in vancomycin-resistant Enterococcus faecium in Germany: Data from the german antimicrobial resistance surveillance (ars). Antimicrob Resist Infect Control 8, 1–11 (2019).

18. Remschmidt, C. et al. Continuous increase of vancomycin resistance in enterococci causing nosocomial infections in Germany -10 years of surveillance. Antimicrob Resist Infect Control 7, 1–7 (2018).

19. Krašovec, R. et al. Spontaneous mutation rate is a plastic trait associated with population density across domains of life. PLoS Biol 15, (2017).

20. Sugino, A., Peebles, C. L., Kreuzer, K. N. & Cozzarelli, N. R. Mechanism of action of nalidixic acid: purification of Escherichia coli nalA gene product and its relationship to DNA gyrase and a novel nicking-closing enzyme. Proc Natl Acad Sci U S A 74, 4767–4771 (1977).

21. Enne, V. I., Delsol, A. A., Roe, J. M. & Bennett, P. M. Rifampicin resistance and its fitness cost in Enterococcus faecium. Journal of Antimicrobial Chemotherapy 53, 203–207 (2004).

22. stp multidrug resistance protein [Mycobacterium tuberculosis H37Rv] - Gene - NCBI. https://www.ncbi.nlm.nih.gov/gene/887274.

23. pyrF - Orotidine 5’-phosphate decarboxylase - Escherichia coli (strain K12) | UniProtKB | UniProt. https://www.uniprot.org/uniprotkb/P08244/entry.

24. Weiss, C. A. et al. NrnA is a 5′-3′ exonuclease that processes short RNA substrates in vivo and in vitro. Nucleic Acids Res 50, 12369 (2022).

25. Chambert, R., Pereira, Y. & Petit-Glatron, M. F. Purification and Characterization of YfkN, a Trifunctional Nucleotide Phosphoesterase Secreted by Bacillus Subtilis. The Journal of Biochemistry 134, 655–660 (2003).

26. De Maat, V., Arredondo-Alonso, S., Willems, R. J. L. & Van Schaik, W. Conditionally essential genes for survival during starvation in Enterococcus faecium E745. BMC Genomics 21, (2020).

27. Jiricny, N. et al. Loss of social behaviours in populations of Pseudomonas aeruginosa infecting lungs of patients with cystic fibrosis. PLoS One 9, (2014).

28. Diggle, S. P., Crusz, S. A. & Cámara, M. Quorum sensing. Current Biology 17, R907–R910 (2007).

29. Muller, M. & Merrett, N. D. Pyocyanin Production by Pseudomonas aeruginosa Confers Resistance to Ionic Silver. Antimicrob Agents Chemother 58, 5492 (2014).

30. Greene, N. P., Kaplan, E., Crow, A. & Koronakis, V. Antibiotic Resistance Mediated by the MacB ABC Transporter Family: A Structural and Functional Perspective. Front Microbiol 9, 950 (2018).

31. Zhu, L. et al. Combined mutations of the penA with ftsX genes contribute to ceftriaxone resistance in Neisseria gonorrhoeae and peptide nucleic acids targeting these genes reverse ceftriaxone resistance. J Glob Antimicrob Resist 35, 19–25 (2023).

32. Bank, C., Ewing, G. B., Ferrer-Admettla, A., Foll, M. & Jensen, J. D. Thinking too positive? Revisiting current methods of population genetic selection inference. Trends in Genetics 30, 540–546 (2014).

33. Bataillon, T. & Bailey, S. F. Effects of new mutations on fitness: insights from models and data. Ann N Y Acad Sci 1320, 76–92 (2014).

34. Bhattacharyya, S. et al. Efflux-linked accelerated evolution of antibiotic resistance at a population edge. Mol Cell 82, 4368-4385.e6 (2022).

35. Webber, M. A. & Piddock, L. J. V. The importance of efflux pumps in bacterial antibiotic resistance. Journal of Antimicrobial Chemotherapy 51, 9–11 (2003).

36. Shigemura, K. et al. Association of overexpression of efflux pump genes with antibiotic resistance in Pseudomonas aeruginosa strains clinically isolated from urinary tract infection patients. The Journal of Antibiotics 2015 68:9 68, 568–572 (2015).

37. Chen, H., Sapula, S. A., Turnidge, J. & Venter, H. The effect of commonly used non-antibiotic medications on antimicrobial resistance development in Escherichia coli. npj antimicrobials and resistance 3, 73 (2025).

38. Yu, X. H. et al. Increased Expression of Efflux Pump norA Drives the Rapid Evolutionary Trajectory from Tolerance to Resistance against Ciprofloxacin in Staphylococcus aureus. Antimicrob Agents Chemother 66, (2022).

39. Lu, J. et al. Safety assessment of Enterococcus lactis based on comparative genomics and phenotypic analysis. Front Microbiol 14, 1196558 (2023).

40. Zhu, K., Chen, S., Sysoeva, T. A. & You, L. Universal antibiotic tolerance arising from antibiotic-triggered accumulation of pyocyanin in Pseudomonas aeruginosa. PLoS Biol 17, (2019).

41. Dietrich, L. E. P., Teal, T. K., Price-Whelan, A. & Newman, D. K. Redox-active antibiotics control gene expression and community behavior in divergent bacteria. Science (1979) 321, 1203–1206 (2008).

42. Meirelles, L. A., Perry, E. K., Bergkessel, M. & Newman, D. K. Bacterial defenses against a natural antibiotic promote collateral resilience to clinical antibiotics. PLoS Biol 19, e3001093 (2021).

43. Meirelles, L. A. & Newman, D. K. Phenazines and toxoflavin act as interspecies modulators of resilience to diverse antibiotics. Mol Microbiol 117, 1384–1404 (2022).

44. Perry, E. K., Meirelles, L. A. & Newman, D. K. From the soil to the clinic: the impact of microbial secondary metabolites on antibiotic tolerance and resistance. Nature Reviews Microbiology 2021 20:3 20, 129–142 (2021).

45. Zhang, C., Kong, Y., Xiang, Q., Ma, Y. & Guo, Q. Bacterial memory in antibiotic resistance evolution and nanotechnology in evolutionary biology. iScience 26, (2023).

46. Luria, S. & Delbrück, M. Mutations of bacteria from virus sensitivity to virus resistance. Genetics 28, 491–511 (1943).

47. Sambrook, J. & Russell, D. W. Purification of nucleic acids by extraction with phenol:chloroform. CSH Protoc 2006, pdb.prot4455 (2006).

