## Supplementary Information for "Pyocyanin produced by *Pseudomonas aeruginosa* Creates Legacy Effects That Boost Antibiotic Resistance Evolution in Enterococci"

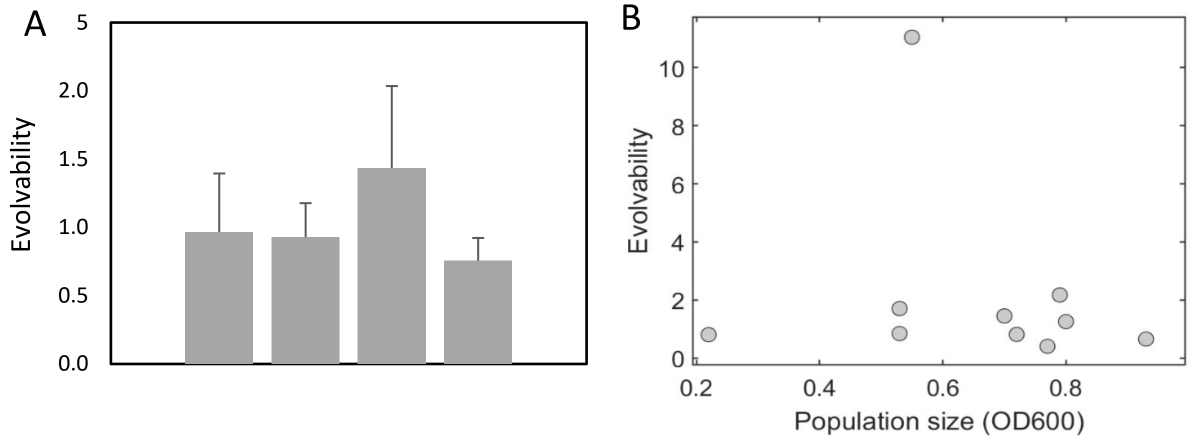

**Figure S1 Evolvability of rifampicin resistance in *E. faecium* is not correlated with the level of nutrients or population size.**

**A Different nutrient level does not lead to changes in evolvability of rifampicin resistance**. Evolvability, as measured by the evolvability assay, of four replicate experiments (each n = 3) under higher nutrient conditions (1.5 x LB) normalized to evolvability under lower nutrient conditions (1 x LB) ($p$ > 0.05, Student's t-test, bar graphs show averages with whiskers depicting the standard deviations). **B No correlation between population size in conditioned medium and evolvability.** Population size versus average rifampicin evolvability of *E. faecium* grown under different conditioned medium conditions. Population size measured after 24 hours and evolvability relative to the evolvability in LB: *K. pneumoniae*, *E. coli* (two different isolates), *P. mirabilis* (two different isolates), *S. aureus*, *S. haemolyticus*, *E. faecium*, *E. faecalis*, and *P. aeruginosa*. Population size in the presence of conditioned medium does not correlate with evolvability of rifampicin resistance due to exposure to that conditioned medium (Materials and Methods). Only the evolvability of rifampicin resistance of *E. faecium* due to the exposure to *P. aeruginosa* conditioned medium is markedly higher (Evolvability > 10).

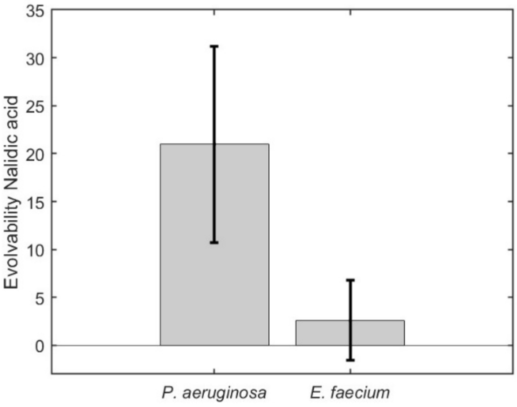

**Figure S2 Increased evolvability of nalidixic acid antibiotic resistance after *P. aeruginosa* exposure.**

**Increased evolvability of nalidixic resistance of *E. faecium* under the influence of *P. aeruginosa* conditioned medium.** Evolvability assay with nalidixic acid instead of rifampicin (Materials and Methods) showed that *P. aeruginosa* conditioned medium also leads to an increased nalidixic acid resistance evolvability, whereas conditioned medium of *E. faecium* does not (one-way ANOVA, $p$ = 0.0005, Dunnett's post-hoc test, $p$ = 0.0005 for *P. aeruginosa* conditioned medium, $p$=0.96 for *E. faecium* conditioned medium). Bar graphs show average evolvability with standard deviation.

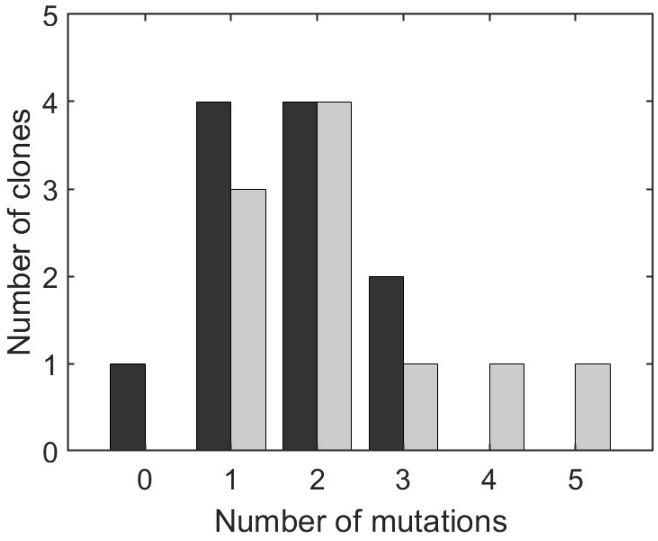

**Figure S3. Distribution of the number of mutations in the resistant clones generated after exposure to *P. aeruginosa* conditioned medium.**

The number of mutations in rifampicin resistant clones generated in the evolvability assay. We sequenced ten rifampicin resistant clones of *E. faecium* generated after transient exposure to *P. aeruginosa* conditioned medium (light bars) and eleven rifampicin resistant clones of *E. faecium* generated after transient exposure to LB controls (black bars). Clones exposed overnight to *P. aeruginosa* conditioned medium have on average more mutations, compared to LB controls (black bars), yet the distributions are not statistically significant due to the limited number of samples (students t-test $p = 0.09$).

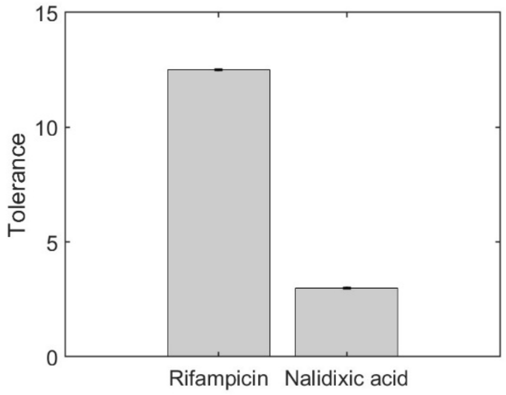

**Figure S4 *P. aeruginosa* conditioned medium mediates antibiotic tolerance.**

*P. aeruginosa* conditioned medium induces antibiotic tolerance of rifampicin and nalidixic acid in *E. faecium*. Bar graphs show the fold increase in tolerance compared to LB reference medium (Materials and Methods), with error bars depicting the standard deviation of the measurement (n = 3).

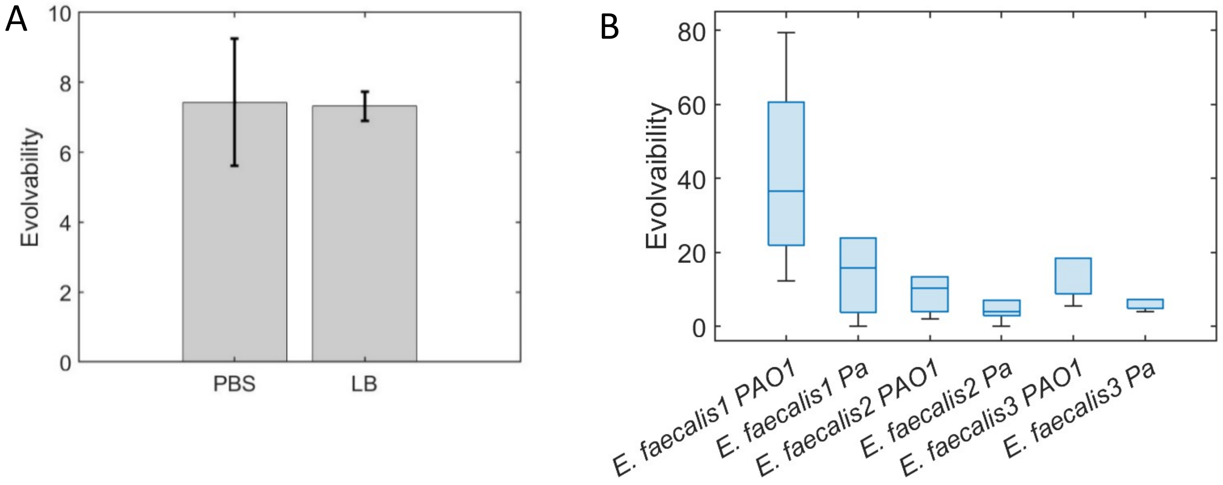

**Figure S5 A The legacy evolvability effect due to *P. aeruginosa*-conditioned medium overnight exposure is conserved after washing in PBS and regrowth in LB.** Washing with PBS, and washing followed by regrowth in LB for three hours maintains the rifampicin resistance evolvability effect due to *P. aeruginosa* conditioned medium (Materials and Methods, Student's t-test, $p > 0.05$). Bar graphs show average evolvability, with standard deviation. **B General effect of exposure to *P. aeruginosa* conditioned medium on antibiotic resistance evolvability of enterococci.** The evolvability of three different *E. faecalis* isolates (1, 2, 3) exposed to PAO1 (canonical *P. aeruginosa*) and uropathogenic *P. aeruginosa* (Pa, used for all experiments described herein) conditioned medium, as measured via the rifampicin evolvability assay (Materials and Methods). The three *E. faecalis* isolates experienced a higher evolvability after the transient exposure to PAO1, compared to the exposure to uropathogenic *P. aeruginosa* medium. The lines in the box plots represent the median, bottom and top edges represent the lower and upper quartiles, whiskers the end points of the dataset.

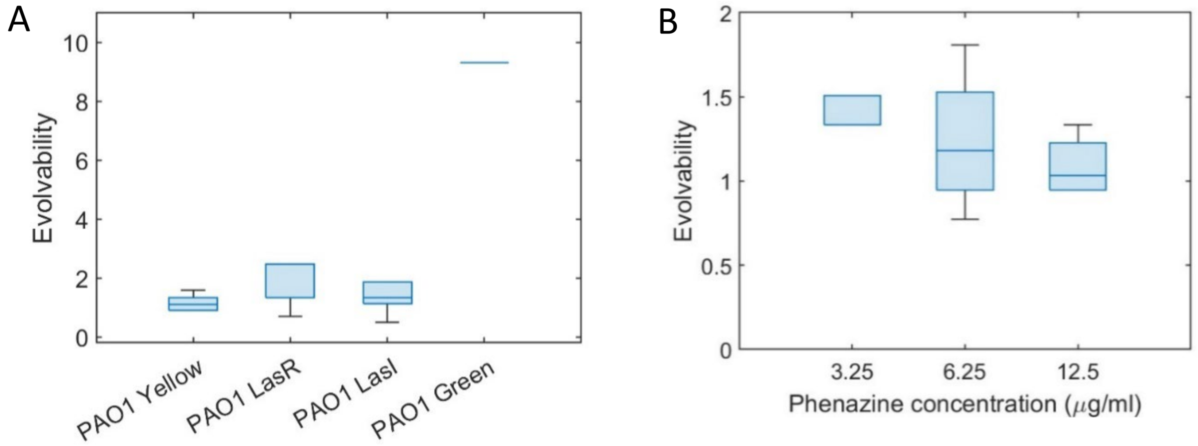

**Figure S6A Increase in antibiotic resistance evolvability associated with quorum sensing of *P. aeruginosa.* A Increase of evolvability of antibiotic resistance by green-tinted version of PAO1 conditioned medium, not by quorum sensing mutants.** Particularly the exposure to the conditioned medium of the aerobically incubated green-tinted version of PAO1 leads to the increase in evolvability, but also the LasR-PAO1-derived quorum sensing mutant grown under lower oxic environments, whereas conditioned medium derived from the un-tinted generated (LB-yellow) PAO1 and LasI PAO1-derived quorum sensing mutant of cultures grown under lower oxic environments do not lead to an increase in evolvability (Materials and Methods, One-way ANOVA, Dunnett's test, PAO1 Yellow $p = 0.9$, PAO1 LasR $p = 0.02$, PAO1 LasI $p = 0.9$, PAO1 Green $p << 0.01$). **B Phenazine does not lead to increased evolvability.** Phenazine, a structural analogue and precursor of pyocyanin does not lead to an increased evolvability of rifampicin antibiotic resistance, as tested via the evolvability assay (Materials and Methods, One-way ANOVA, $p > 0.05$). The lines in the box plots represent the median, bottom and top edges represent the lower and upper quartiles, whiskers the end points of the dataset.

**Figure S7 Differential gene expression of *E .faecium* in the presence of *P. aeruginosa* conditioned medium compared to in *E. faecium* conditioned medium.**

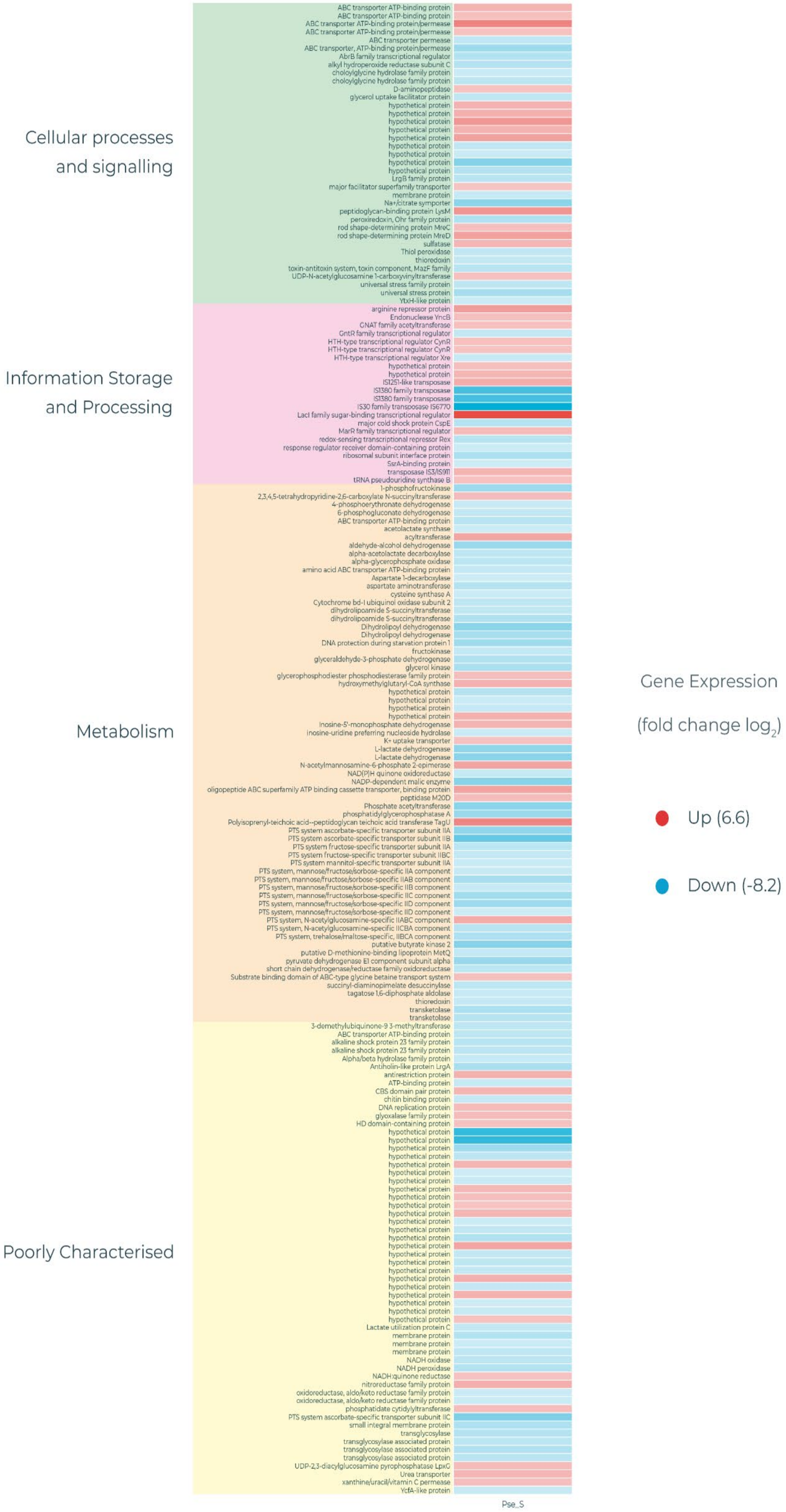

**Table S1**

**Different RpoB mutations generated under the transient exposure of *P. aeruginosa* conditioned medium, compared to the classical RpoB mutations in LB medium.**

| **Conditioned medium *P. aeruginosa*** | **LB** |
|---|---|
| Q473K (CAG→AAG) | S491L (TCA→TTA) |
| Q473K (CAG→AAG) | S491L (TCA→TTA) |
| H486L (CAT→CTT) | S491L (TCA→TTA) |
| H486D (CAT→GAT) | H486Y (CAT→TAT) |
| G482D (GGT→GAT) | H486Y (CAT→TAT) |
| | H486Y (CAT→TAT) |
| | H486Y (CAT→TAT) |
| | H486Y (CAT→TAT) |
| | G482D (GGT→GAT) |

RpoB mutations of rifampicin resistant clones derived from the phenotypic evolvability assay, after overnight exposure to LB or *P. aeruginosa* conditioned medium. After overnight exposure to LB the 'classical' RpoB mutations are found in rifampicin resistant clones (Enne et al., 2004). After overnight exposure to *P. aeruginosa*, different RpoB mutations than the 'classical' RpoB mutations are found in rifampicin resistant clones. Generally, fewer RpoB mutations were found in clones that were previously exposed to *P.aeruginosa*, compared to those exposed to LB.

**Table S2**

**Mutations other than RpoB in the mutants derived from evolvability assay**

Other mutations than RpoB in rifampicin resistant clones derived from the phenotypic evolvability assay, after overnight exposure to LB or *P. aeruginosa* conditioned medium. After overnight exposure, more other mutations were found in clones that were previously exposed to *P. aeruginosa*, compared to those exposed to LB. The latter mostly had the 'classical' RpoB mutations (Enne et al., 2004).

| ***P. aeruginosa* conditioned medium** | | **LB reference medium** | |
|---|---|---|---|
| *pdhD* | Dihydrolipoyl dehydrogenase | *L_02668 →* | hypothetical protein |
| *ftsA_1* | Cell division protein FtsA | *yfkN_2 ←* | Trifunctional nucleotide phosphoesterase protein YfkN |
| *yfkN_2 ← / ← glmU* | Trifunctional nucleotide phosphoesterase protein YfkN/Bifunctional protein GlmU | *eutB ←* | Ethanolamine ammonia-lyase heavy chain |
| *pyrF* | Orotidine 5'-phosphate decarboxylase | *lytR_1 ←* | Transcriptional regulator LytR |
| *nrnA* | putative bifunctional oligoribonuclease and PAP phosphatase NrnA (2 clones) | *pbpF_1 ←* | Penicillin-binding protein 1F |
| *carA* | Carbamoyl-phosphate synthase small chain | *L_03056 →* | hypothetical protein |
| *sugC_1 ← / → mgsA* | Trehalose import ATP-binding protein SugC/Methylglyoxal synthase | *L_01950 →* | hypothetical protein |
| *L_02631* | putative ABC transporter ATP-binding protein | *L_00734 →* | hypothetical protein |
| *helD ← / → ycfH* | Helicase IV/putative metal-dependent hydrolase YcfH (2 clones) | | |
| *stp* | Multidrug resistance protein Stp | | |
| *yfkN_2* | Trifunctional nucleotide phosphoesterase protein YfkN | | |

| | |
|---|---|
| *L_00636* ← / → *L_00637* | hypothetical protein/hypothetical protein |
| *L_03056* → | hypothetical protein |
| *L_02442* ← | hypothetical protein |

**Table S3 List of the ten most overexpressed genes of *E. faecium* exposed to *P. aeruginosa* conditioned medium versus *E. faecium* exposed to *E. faecium* conditioned medium, assessed via transcriptomics.**

| | Log2-fold change | FDR | p-value |
|---|---|---|---|
| LacI family sugar-binding transcriptional regulator | 4.17 | 1.04e-8 | 1.02e-11 |
| Polyisoprenyl-teichoic acid--peptidoglycan teichoic acid transferase TagU | 2.95 | 2.38e-6 | 1.40e-8 |
| ABC transporter ATP-binding protein/permease MacB FtsX-like permease family | 3.01 | 1.64-e5 | 12.4e-7 |
| Rod shape-determining protein MreD | 2.31 | 1.24e-4 | 1.33e-6 |
| Oligopeptide ABC superfamily ATP binding cassette transporter, binding protein OppA | 2.28 | 1.91e-4 | 2.44e-6 |
| ABC transporter ATP-binding protein | 1.91 | 9.84e-4 | 2.32e-5 |
| Peptidoglycan-binding protein LysM | 2.61 | 1.48e-5 | 1.06e-7 |
| PTS system, N-acetylglucosamine-specific IIABC component ptsG PTS_EIIB,PTS_EIIC | 2.09 | 1.94e-3 | 6.37e-5 |
| ABC transporter ATP-binding protein | 1.61 | 2.53e-3 | 9.15e-5 |
| CBS domain pair protein YkuL | 1.86 | 1.90e-3 | 5.91e-5 |
| MarR family transcriptional regulator | 1.72 | 1.81e-3 | 5.39e-5 |
